# The IRE1α stress signaling axis is a key regulator of neutrophil antimicrobial effector function

**DOI:** 10.1101/743336

**Authors:** B. H. Abuaita, G. J. Sule, T. L. Schultz, F. Gao, J. S. Knight, M. X. O’Riordan

## Abstract

Activation of the endoplasmic reticulum stress sensor, IRE1α, is required for effective immune responses against bacterial infection and is associated with human inflammatory diseases where neutrophils are a key immune component. However, the specific role of IRE1α in regulating neutrophil effector function has not been studied. Here we show that infection-induced IRE1α activation licenses neutrophil antimicrobial capacity, including IL-1β production, NET formation, and MRSA killing. Inhibition of IRE1α diminished production of mitochondrial reactive oxygen species (mROS) and decreased CASPASE-2 activation, which both contributed to neutrophil antimicrobial activity. Mice deficient in Caspase-2 were highly susceptible to MRSA infection and failed to form NETs in a subcutaneous abscess. IRE1α activation enhanced calcium influx and citrullination of histone H3 (Cit-H3) independently of mROS production, suggesting that IRE1α coordinates multiple pathways required for NET formation. Our data demonstrate that the IRE1α-Caspase-2 axis is a major driver of neutrophil activity against MRSA infection and highlight the importance of IRE1α in neutrophil antibacterial function.

**One Sentence Summary:** IRE1α controls neutrophil antimicrobial defenses

## Introduction

Neutrophils are critical first responders to infection, poised to rapidly kill invading pathogens with a dense array of secretory granules containing a diverse repertoire of antimicrobial molecules (*1, 2*). Neutrophils can kill intracellular microbes by concentrating these microbicides in phagosomes through fusion of secretory granules and generation of reactive oxygen species (ROS). These polymorphonuclear (PMN) leukocytes can also kill extracellular microbes by degranulation and exocytosis of secretory granules, releasing antimicrobial molecules into the extracellular space (*3*). Additionally, neutrophils secrete inflammatory chemokines and cytokines to recruit other immune cells to mediate a multi-pronged host defense against infection (*4*).

Upon stimulation, neutrophils can release sticky web-like structures composed of genomic DNA and associated proteins, termed neutrophil extracellular traps (NETs) (*5*). NET formation is induced by a wide range of microbes, including bacteria, fungi, parasites and viruses (*6*). During infection, neutrophils extrude NETs to trap microbes, likely limiting spread from the primary site of infection (*7, 8*). Preventing NET formation decreases the capacity of neutrophils to kill Gram-positive and Gram-negative bacteria (*5*). NETs are enriched with neutrophil-derived granule enzymes like neutrophil elastase, myeloperoxidase, cathepsin G, and cathelicidins, defensins and other antimicrobial peptides (*9*). Thus, NETs may aid in microbial killing directly by concentrating microbicidal molecules onto extracellular pathogens trapped in this structure. NETs are also linked to inflammatory diseases such as Systemic Lupus Erythematosus (SLE), psoriasis, or diabetes (*10, 11*). Exaggerated NET formation or defects in NET clearance can exacerbate these diseases (*10*). Thus, NET formation must be highly regulated to promote host defense without resulting in pathological inflammatory consequences.

NET release requires generation of reactive oxygen species (ROS) and histone modification to promote decondensation of nuclear chromatin (*12, 13*). Several chemical and biological stimuli trigger ROS production and histone citrullination to promote NET formation, including phorbol 12-myristate 13-acetate (PMA), the calcium ionophore (A23187) and various microbial infections (*6, 14, 15*). Notably, ROS that contribute to NET formation can be generated by multiple sources including the major phagocyte oxidase, NADPH oxidase 2 (NOX2) and mitochondria (*14, 16, 17*). ROS induce translocation of granule proteins such as neutrophil elastase (NE) into the nucleus to promote chromatin relaxation by cleaving nucleosomal histones (*18*). Concomitantly, histone citrullination by peptidylarginine deiminase-4 (PAD4) reduces net positive charge and decreases histone binding affinity for nuclear DNA, freeing DNA to be expelled from the cell (*13, 19–21*). PAD4 requires calcium as a cofactor to become active, and localizes to the neutrophil nucleus upon stimulation (*22, 23*). Treating neutrophils with PAD4 inhibitors or preventing calcium mobilization suppresses NET formation (*19, 24*). PAD4-deficient neutrophils are unable to citrullinate histones, decondense chromatin and generate NETs, resulting in greater susceptibility to bacterial infection (*20*).

Neutrophils, as highly secretory cells, are dependent on optimal ER function. In general, perturbation of ER homeostasis triggers an adaptive stress response, termed the unfolded protein response (UPR), and is controlled by three resident ER sensors, of which the inositol-requiring enzyme 1-α (IRE1α) is the most evolutionarily conserved (*25*). During ER stress, oligomerization of IRE1α triggers autophosphorylation and activation of its cytoplasmic endonuclease domain (*26, 27*). The endonuclease removes a 26-nucleotide intron from the cytoplasmic *Xbp1* mRNA, licensing translation of the transcription factor X-box Binding Protein-1 (XBP1) (*28*). XBP1 then induces expression of many genes that improve ER protein folding and degradation to restore ER homeostasis (*29, 30*). Activation of IRE1α in immune cells enhances production of many proinflammatory cytokines, such as IL-1β, IL-6, and TNFα (*31–33*). IRE1α is implicated in inflammation during infection and in many autoimmune diseases such as rheumatoid arthritis and systemic lupus erythematosus, where neutrophils and NET formation contribute to the progression of these diseases (*10, 34*).

We recently demonstrated that IRE1α aids in clearance of methicillin-resistant *Staphylococcus aureus* (MRSA) in a subcutaneous abscess model, normally controlled by neutrophils and neutrophil-derived IL-1β (*35, 36*). However, the specific role of IRE1α in neutrophil function is poorly understood. Here we investigated the contribution of IRE1α to neutrophil antimicrobial function. We show that MRSA infection triggers neutrophil IRE1α activation, and find that this activation is required for efficient neutrophil killing of MRSA, production of IL-1β and NET formation. IRE1α promotes generation of mitochondrial ROS (mROS), activation of CASPASE-2 and intracellular calcium mobilization, all required for optimal neutrophil antimicrobial function. Moreover, Caspase-2 deficient mice infected with MRSA exhibited increased bacterial burden, lower levels of IL-1β, and failed to generate NETs in a subcutaneous abscess model compared to wild-type mice. Collectively, these data point to IRE1α as a positive regulator of neutrophil effector function, which is critical in coordinating the inflammatory response.

## Results

### Neutrophil IRE1α activation enhances MRSA killing and IL-1β production

To define the role of IRE1α in neutrophil host defense against bacterial infection, we first determined whether infection activates neutrophil IRE1α, using the clinically derived MRSA strain USA300 LAC. Human PMN were purified from blood of healthy volunteers as previously described (*37*), yielding cell populations >95% CD15^+^CD16^+^, characteristic of neutrophils (Fig. S1A). Neutrophils incubated with MRSA showed increased levels of the spliced variant of *XBP1*, indicating that IRE1α was activated (Fig. 1A and B). To determine the functional consequences of MRSA-induced IRE1α activation on neutrophil bactericidal activity and IL-1β production, we treated PMN with IRE1α inhibitors 4μ8C or KIRA6 (*38, 39*). In inhibitor-treated neutrophils, *XBP1* splicing was substantially reduced (Fig. 1B), and these cells were less capable of killing MRSA (Fig. 1C). In addition, inhibitor-treated neutrophils produced lower levels of IL-1β (Fig. 1D) compared to controls. To determine whether these outcomes were due to drug toxicity on MRSA and/or neutrophils, we monitored effects of 4μ8C or KIRA6 on MRSA growth and neutrophil survival. Treatment with 4μ8C or KIRA6 did not affect either axenic growth of MRSA (Fig. S1, B and C) or neutrophil cell viability (Fig. S1, D and E). These data demonstrate the requirement for IRE1α in neutrophil killing and IL-1β production upon MRSA infection.

**Fig. 1.**
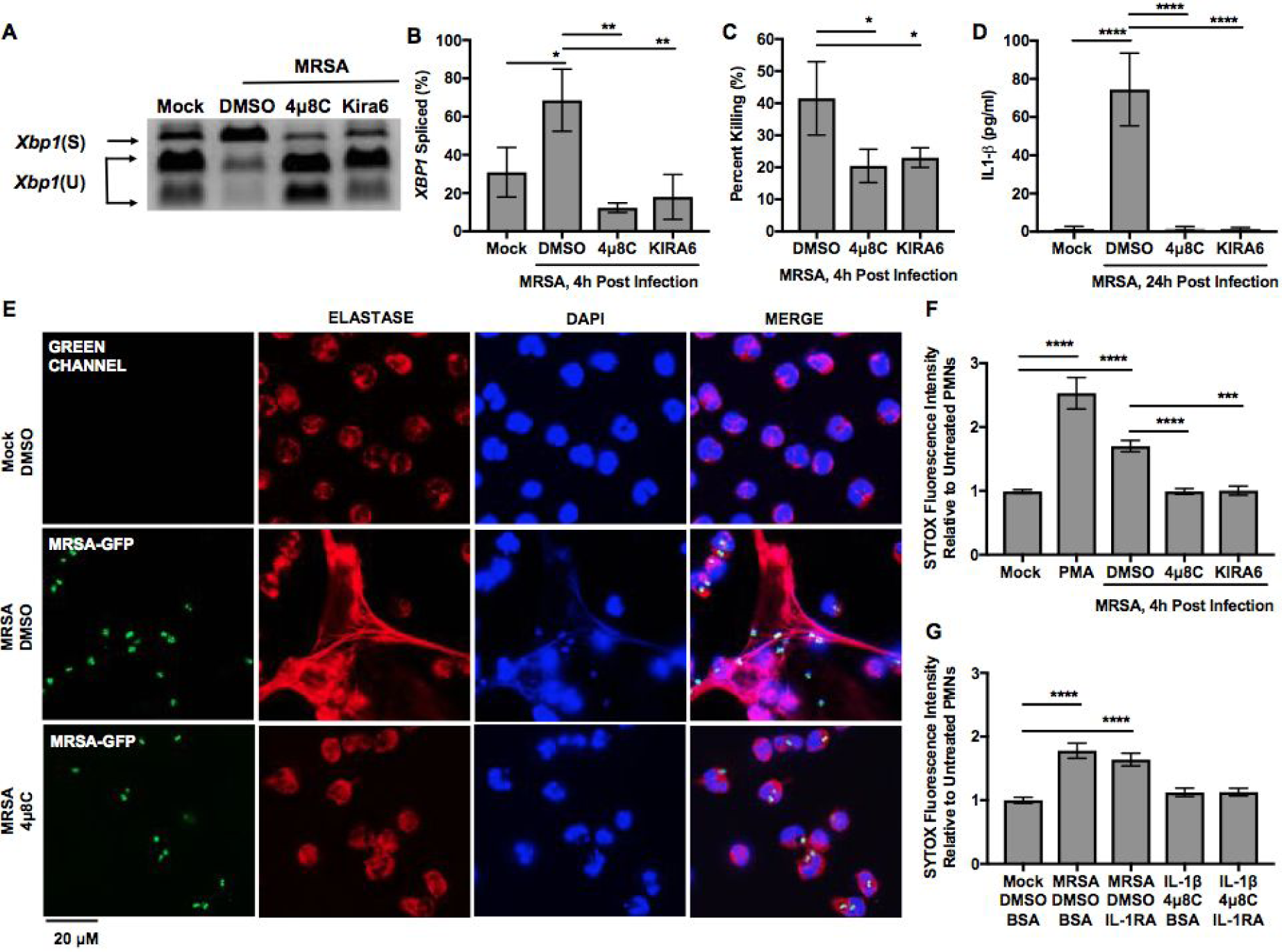
Human neutrophil IRE1α is required for bacterial killing, IL-1β production and NET formation. **(A)** *XBP1* splicing was assessed at 4hour post MRSA-infection (4h pi) by RT-PCR, followed by *PstI* digestion to cleave the unspliced product. Because unspliced mRNA contains a *PstI* site within the 26 bp intron, the digested PCR products yield two smaller fragments representing unspliced (U) *XBP1* and one larger fragment representing spliced (S) *XBP1.* (**B)** Percent of spliced *XBP1* was quantified using ImageJ based on band intensity. **(C)** IL-1β in culture medium was measured by ELISA at 24h pi. **(D)** Percent killing was quantified by percent difference in CFU at 4h in the presence of neutrophils relative to bacteria cultured alone. **(E)** Representative fluorescence microscopy images of human neutrophils left untreated (Mock) or infected with MRSA-GFP (Green) for 4h ±IRE1α inhibitor, 4*µ*8C. Cells were labeled with anti-Human Elastase antibody (Red), stained with DAPI (Blue) to label DNA, and imaged using epifluorescence microscopy. **(F)** NET formation was quantified by cell-impermeant SYTOX Green nucleic acid dye. Neutrophils were cultured with 500 nM SYTOX Green and fluorescence intensity measured at 4h. Cells were left untreated or infected with MRSA in presence of 25 µM 4*µ*8C, 10 µM KIRA6 or DMSO. (**G)** Quantification of NET formation by SYTOX Green plate reader assay when neutrophils left untreated, and stimulated with IL-1β or infected with MRSA in the absence or presence excess interleukin-1 receptor antagonist IL-1RA. Graphs indicate mean +/-SD except SYTOX Green graphs indicate mean +/- SEM of n≥3 independent experiments. P value was calculated using one-way ANOVA with post-Tukey’s test for multiple comparisons. p value: *< 0.05, **< 0.01, ***< 0.001 and ****<0.0001.

### NET formation depends on IRE1α activity during MRSA infection

Neutrophils form bactericidal extracellular traps (NETs) during *Staphylococcus aureus* infection (*5, 8*). NET formation is also driven by IL-1β production and signaling (*40, 41*). Therefore, we investigated the role of IRE1α in NET formation stimulated by MRSA infection. Neutrophils were cultured on poly-lysine coated coverslips and infected with MRSA-GFP at a multiplicity of infection (MOI) of 10 for 4 hours in the presence and absence of IRE1α inhibitors. NET formation was assessed by immunofluorescence microscopy by measuring the extracellular co-localization of neutrophil elastase (NE) and DNA (Fig. 1E). Uninfected neutrophils predominantly had a multi-lobed nucleus and intracellular staining of NE, indicating that these cells did not form NETs. During MRSA infection, NE and DNA were observed extracellularly in many cells, while others lost the multi-lobed nucleus appearance, suggesting that these cells had undergone NETosis. When neutrophils were treated with IRE1α inhibitor 4μ8C during MRSA infection, we observed lower levels of extracellular NE and DNA association compared to the DMSO control, indicating that IRE1α contributes to MRSA-induced NETosis. To quantify NETosis, we measured accumulation of extracellular DNA by using the cell-impermeable dye SYTOX Green, which fluoresces upon binding to nucleic acids. MRSA infection stimulated neutrophils to release extracellular DNA, indicated by increased SYTOX fluorescence intensity compared to untreated neutrophils (Fig. 1F). Similar to the fluorescence microscopy data, treatment with IRE1α inhibitors reduced SYTOX fluorescence intensity during MRSA infection to a level similar to untreated neutrophils, indicating that the IRE1α endonuclease activity is required for NET formation during MRSA infection.

IL-1β signaling is required for NETosis during sterile inflammatory acute gout (*40*). We therefore investigated whether IRE1α controls MRSA-induced NET formation via IL-1β signaling. Treatment of neutrophils with IL-1 receptor antagonist, IL-1RA, prior to MRSA infection did not decrease NET production, which was quantified by extracellular SYTOX Green fluorescence (Fig. 1G). Neutrophils stimulated with recombinant IL-1β in the presence or absence of IL-1RA and 4μ8C (Fig. 1G). Treatment of uninfected neutrophils with recombinant IL-1β, regardless of the presence of IL-1RA or 4μ8C, did not increase SYTOX fluorescence intensity compared to untreated neutrophils, indicating that IL-1β is not sufficient to cause neutrophils derived from healthy donors to undergo NETosis. Together, our data suggest that IRE1α promotes neutrophil NETosis in response to MRSA infection independently of IL-1β signaling.

### Mitochondrial ROS contributes to IRE1α-dependent NET formation

NET formation occurs in an oxidant-dependent manner. NADPH oxidase 2 (NOX2) is a major complex for generation of reactive oxygen species (ROS) and is required for some stimuli-induced NET formation (*16*). However, ROS production by mitochondria (mROS) is also important for NET formation during sterile inflammation and has been associated with NOX2-independent NETosis (*14, 17*). We recently showed that macrophage IRE1α induces mROS in response to MRSA infection, which is required for MRSA killing in phagosomes (*42*). Thus, we hypothesized that neutrophil IRE1α controls NET formation during MRSA infection via production of mROS. To test this hypothesis, we first monitored production of total ROS and mROS by neutrophils during MRSA infection. Total ROS was measured by flow cytometry using CM-H_2_DCFDA probe, which detects many cellular free radical species, and mitochondrial hydrogen peroxide (mH_2_O_2_), measured by the MitoPY1 probe (*42–44*). MRSA infection increased both total and mH_2_O_2_ at 4h post-infection (pi) (Fig. 2, A and B). When neutrophils were treated with IRE1α inhibitors or the mROS specific scavenger (NecroX-5), total ROS and mH_2_O_2_ production was decreased. Notably, mH_2_O_2_ production was profoundly decreased by IRE1α inhibition when compared to total ROS, suggesting that a decrease in total ROS was likely due to decreased mH_2_O_2_ production. In addition to increased mH_2_O_2_ production, MRSA infection caused reduction in mitochondrial membrane potential, which was rescued by treatment with IRE1α inhibitors or the mROS scavenger, NecroX-5, prior to MRSA infection (Fig. 2C). We next investigated the role of mROS production in the ability of neutrophils to kill MRSA and form NETs. Treating neutrophils with the mROS scavenger prior to MRSA infection decreased MRSA killing (Fig. 2D) and suppressed NET formation (Fig. 2, E and F). Thus, our data point to a role for IRE1α activation in promoting neutrophil mH_2_O_2_ production, which is required for MRSA-induced NETosis and bactericidal function. Mitochondria-derived molecules such as ATP, mROS, Cardiolipin, Cytochrome c and oxidized DNA are potent modulators of innate immune pathways (*45*). We recently found that the IRE1α-Caspase-2 axis mediates NLRP3 inflammasome activation in *Brucella-*infected macrophages (*32*). Caspase-2 acted upstream of mitochondrial damage, coordinating release of mitochondrial components into the cytosol to initiate activation of the inflammasome. We therefore investigated whether CASPASE-2 played a role in mitochondrial damage in neutrophils and NETosis during MRSA infection. We monitored CASPASE-2 activation in neutrophils upon MRSA infection using the fluorescent probe FLICA FAM-VDVAD that covalently binds to active CASPASE-2. MRSA infection activated CASPASE-2 in 64% of cells (Fig. 3A). Pretreatment of cells with CASPASE-2 inhibitor (z-VDVAD), 4μ8C and NecroX-5 prior to infection prevented CASPASE-2 activation, suggesting that activity of IRE1α and mROS production is necessary for CASPASE-2 activation in neutrophils and indicates that the activation of IRE1α-Caspase-2 axis is conserved between neutrophils and macrophages in response to microbial infection (*32*). Importantly, CASPASE-2 inhibition alleviated the loss of mitochondrial membrane potential (Fig. 3B), decreased NET formation (Fig. 3, C and D) and reduced IL-1β production (Fig. 3E) during MRSA infection. Caspase-2 has previously been implicated in promoting apoptosis through activation of Caspase-3 (*46, 47*). A recent study showed that CASPASE-3 mediated apoptosis and NETosis occur concurrently under certain conditions, such as exposing neutrophils to high UV radiation (*48*). To determine whether CASPASE-3 dependent apoptosis occurs during MRSA infection and whether the IRE1α-Caspase-2 axis plays a role in this process, we monitored CASPASE-3 activity during MRSA infection when IRE1α and CASPASE-2 are inhibited (Fig. 3F).

**Fig. 2.**
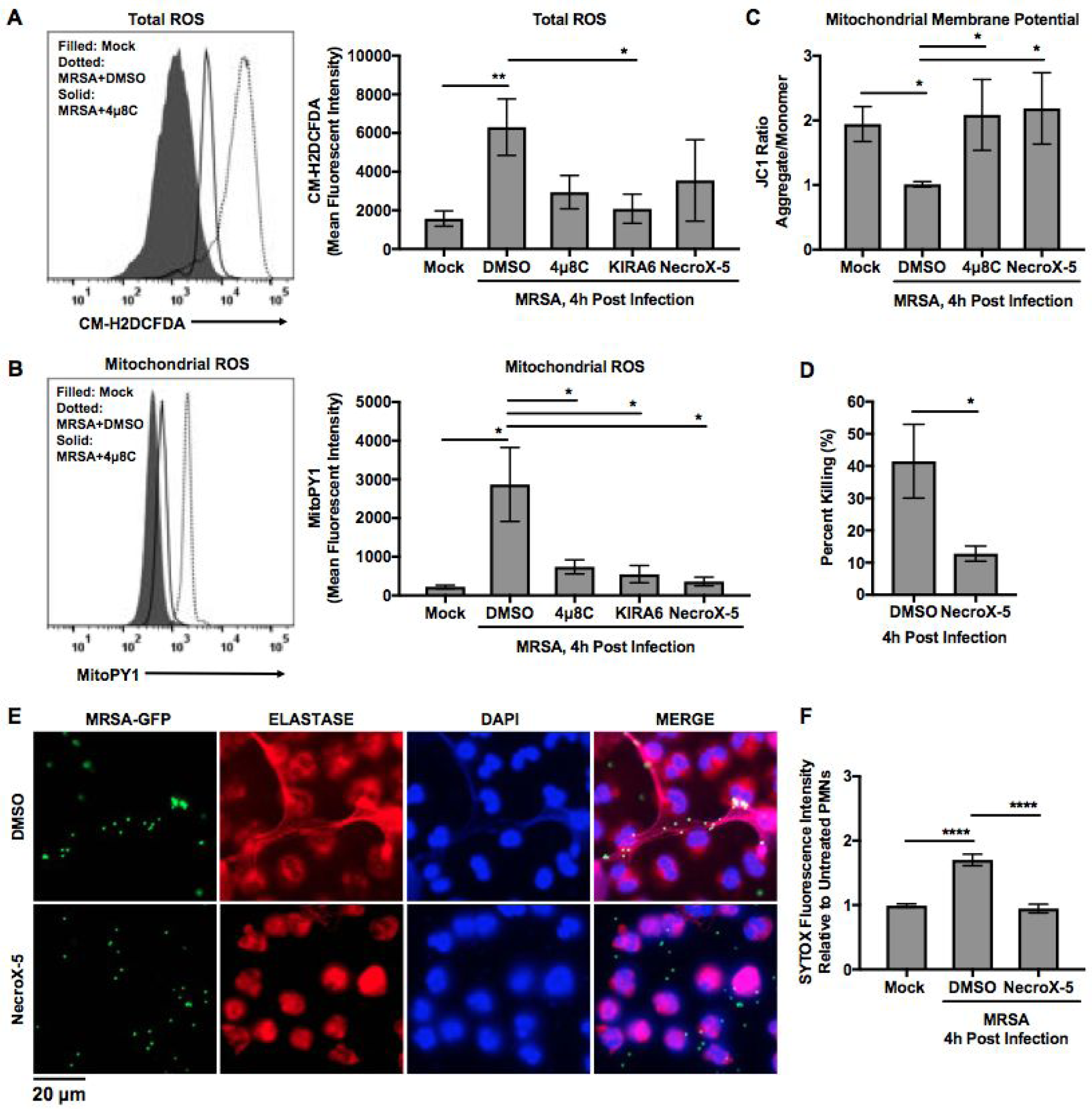
IRE1α controls neutrophil antimicrobial function via mROS production. **(A)** Total cellular ROS production was assessed by flow cytometry at 4 hpi using CM-H_2_DCFDA dye. (Right) Quantification of mean fluorescence intensity (MFI) under the indicated conditions. **(B)** Mitochondrial ROS production was monitored by flow cytometry using the mitochondria-targeted probe, MitoPY1. (Right) Mean fluorescence intensity (MFI) of neutrophils after 1 hr labeling with MitoPY1 followed by MRSA infection in the presence of indicated inhibitors and analyzed at 4 hpi. **(C)** Mitochondrial membrane potential was assessed by flow cytometry at 4 hpi in the presence of indicated inhibitors using JC1 dye. Ratiometric analysis of red fluorescence (FL2) to green fluorescence (FL1) was used to determine the mitochondrial membrane potential status. **(D)** Percent MRSA killing by neutrophils in the presence or absence of NecroX-5. Percent killing was calculated by percent difference in CFU at 4h in the presence of neutrophils relative to bacteria cultured alone. **(E)** NET formation was monitored by fluorescence microscopy and quantified by SYTOX Green assay **(F)** at 4 pi. Graph is presented as the mean of n≥3 independent experiments +/- SEM. MFI quantification was determined using FlowJo software, representing the geometric mean. MFI obtained from unstained cells was subtracted from the MFI of all stained samples. Unless otherwise stated, graphs indicate the mean of n≥3 independent experiments +/- SD. *P* value was calculated using one-way ANOVA with Tukey’s post-test for multiple comparisons. *P* value: *< 0.05, **< 0.01 and ****<0.0001.

**Fig. 3.**
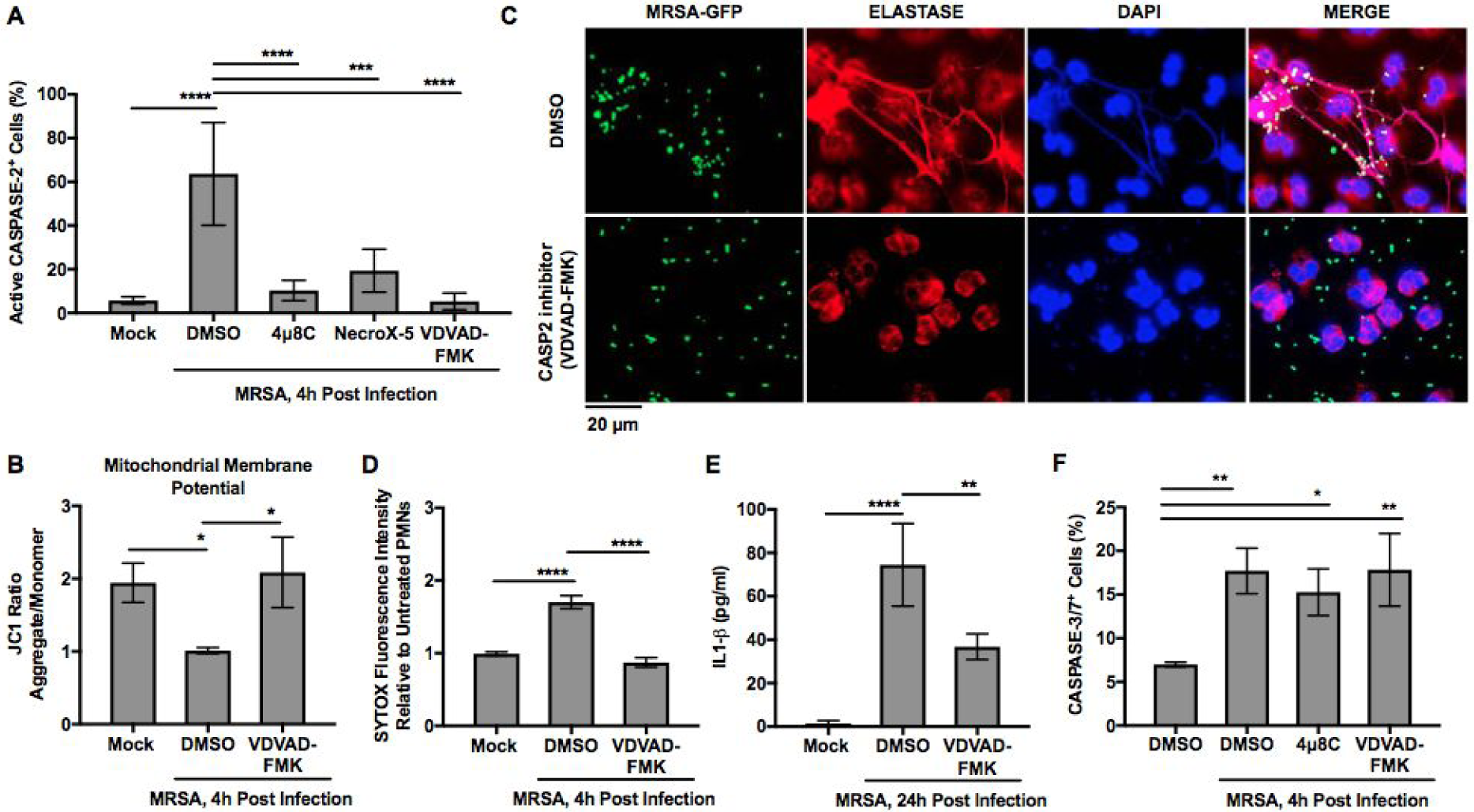
IRE1α-CASPASE-2 axis controls MRSA-induced NET formation and IL1-β production. (A) Percent active CASPASE-2^+^ neutrophils when left untreated (mock) or infected with MRSA in the presence of indicated inhibitors. CASPASE-2 activation was monitored by flow cytometry using the fluorescent probe (FAM-VDVAD-FMK), which irreversibly binds to activated CASPASE-2. Percent CASPASE-2 activated cells was determined by gating against unstained cells. (B) Mitochondrial membrane potential was monitored by flow cytometry using a ratiometric measurement of JC1 dye (FL2/FL1). (C) Representative microscopy images of human neutrophils infected with MRSA-GFP for 4h (green) ±10 µM CASPASE-2 inhibitor VDVAD-FMK, and stained for NE (red) and DNA (blue). (D) Quantification of NET formation by SYTOX Green was performed with neutrophils infected with MRSA for 4h +CASPASE-2 specific inhibitor, VDVAD-FMK or DMSO. Graph shows the mean of n≥3 independent experiments +/- SEM. (E) IL-1β production by human neutrophils when left untreated or infected with MRSA for 24h ± CASPASE-2 inhibitor. (F) CASPASE-3/7 activity of human neutrophils left untreated (mock) or infected with MRSA for 4h (MRSA) ± indicated inhibitors. CASPASE-3/7 activity was measured by flow cytometry (CellEvant CASPASE-3/7 assay). Percent CASPASE-3/7^+^ cells was determined by gating against unstained cells. Unless otherwise stated, graphs indicate means +/- SD of n≥3 independent experiments. *P* value was calculated using one-way ANOVA with Tukey’s post-test for multiple comparisons. *P* value: *< 0.05, **< 0.01, ***< 0.001 and ****<0.0001.

Neutrophils infected with MRSA exhibited CASPASE-3/7 activity. However, treatment with CASPASE-2 or IRE1α inhibitors did not interfere with CASPASE-3 activation, indicating that IRE1α and CASPASE-2 are not involved in MRSA-induced CASPASE-3 activation. These data suggest that the IRE1α-Caspase-2 axis mediates NETosis independently of CASPASE-3 mediated apoptosis during MRSA infection.

### IRE1α augments histone citrullination and calcium influx independently of mROS

Histone modification by citrullination is a vital process for NET formation (*19*). In neutrophils, histone citrullination is carried out by the activity of PAD4, which requires intracellular calcium mobilization (*49*). To test whether IRE1α and mROS contribute to MRSA-induced histone citrullination and calcium influx, we assessed induction of histone citrullination during MRSA infection by immunoblot analysis using an antibody specific for the citrullinated form of histone H3. MRSA infection triggered an increase in histone H3 citrullination (Fig. 4A). Treatment with IRE1α inhibitors prior to infection reduced the level of citrullinated histone H3 compared to untreated neutrophils. However, treatment with the mROS scavenger, NecroX-5, showed similar levels of histone H3 citrullination compared to DMSO-treated cells (Fig. 4B). We then monitor calcium flux in neutrophils during MRSA infection using the calcium fluorescent indicator dye, Fluo-4 AM. The kinetic fluorescence intensity was monitored by flow cytometry on live cells. There was a sharp, transient increase in Fluo-4 AM fluorescence intensity during MRSA infection, indicating a spike in intracellular calcium (Fig. 4C). This increase was followed by a decline in fluorescence intensity that never reached baseline value observed in non-stimulated PMN. Treatment with an IRE1α inhibitor, but not an mROS scavenger, suppressed calcium flux in MRSA-infected cells. Stimulation with a calcium ionophore, A23187, caused a rapid increase in calcium similar to MRSA infection (Fig. 4D). However, the ionophore-induced increase in calcium influx did not decline over time, and treatment with an IRE1α inhibitor or mROS scavenger did not decrease calcium influx. These data indicate that IRE1α is critical for infection-induced calcium mobilization and histone modification and suggest that IRE1α controls NET formation via mROS-dependent and independent mechanisms.

**Fig. 4.**
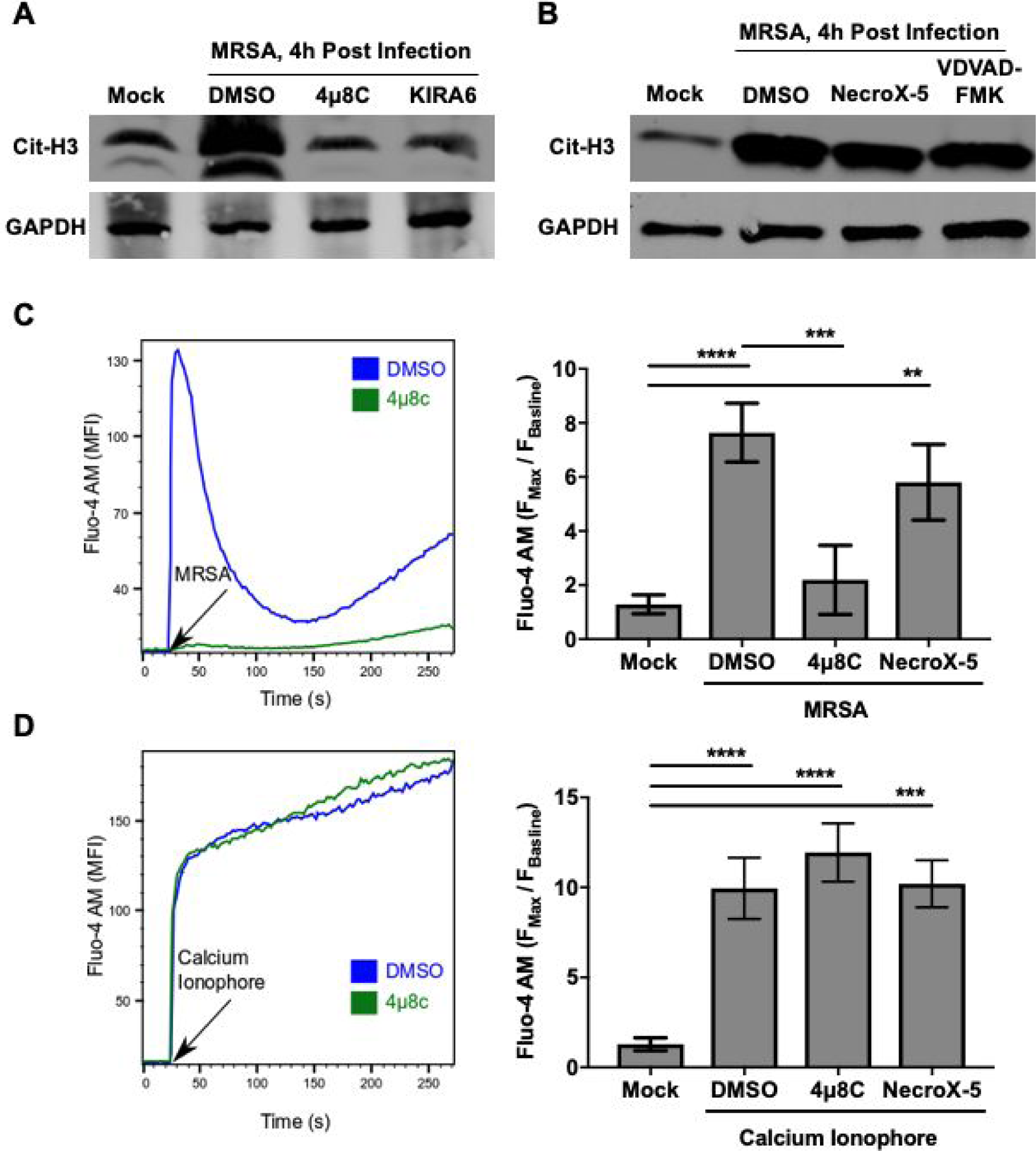
IRE1α activation promotes histone citrullination and calcium influx during MRSA infection. **(A)** Immunoblot analysis of untreated (mock) and MRSA-infected neutrophils (4h) ±IRE1α inhibitors with citrullinated H3 antibody. GAPDH was used as a loading control. **(B)** Immunoblot analysis of citrullinated histone H3 and GAPDH from cell lysates of neutrophils when left untreated (mock) or infected with MRSA for 4h with indicated inhibitors. Fluo-4 AM fluorescence intensity of human neutrophils for calcium influx during MRSA infection **(C)** and calcium ionophore (10 µM A23187) stimulation **(D)** in the presence of IRE1α inhibitor (25 µM 4µ8C), mROS scavenger (10 µM NecroX-5) and DMSO control. Data were acquired by flow cytometry and analyzed by FlowJo software for mean fluorescence intensity of n≥3 independent experiments +/- SD. *P* value was calculated using one-way ANOVA with Tukey’s post-test for multiple comparisons. *P* value: **< 0.01, ***< 0.001 and ****<0.0001.

### Caspase-2 augments NET formation and MRSA clearance in the subcutaneous abscess

Neutrophil IRE1α-Caspase-2 axis can influence host defenses in *vivo* against MRSA infection via generation of antimicrobial NETs. Caspase-2 has been shown to promotes cell death in epithelial cells during *S. aureus* infection (*50*) and in macrophages during *Salmonella* infection (*51*), which may impact host defenses. But, the contribution of Caspase-2 to antimicrobial host defenses has not been tested. To evaluate the role of Caspase-2 during MRSA infection *in vivo*, we used a mouse model of subcutaneous MRSA infection and monitored inflammation, NET formation and bacterial killing. WT and Caspase-2 deficient mice (*Casp2^-/-^*) were infected with MRSA (10^7^ CFU) subcutaneously on the right flank. Bacterial burden and cytokine levels in the abscess were measured at 3d pi. In comparison to WT mice, there were significantly higher bacterial counts (Fig. 5A) and higher TNFα levels in the abscesses of *Casp2^-/-^* mice (Fig. 5B). This increased level of TNFα in *Casp2^-/-^* abscesses may reflect the higher bacterial burden. However, compared to WT mice, *Casp2^-/-^* mice had a decreased level of abscess IL-1β (Fig. 5C), which is critical for clearance of *S. aureus* from a skin and soft-tissue infection model (*36*). Thus, Caspase-2 contributes to IL-1β production and is necessary for *S. aureus* clearance in a mouse subcutaneous abscess model.

**Fig. 5.**
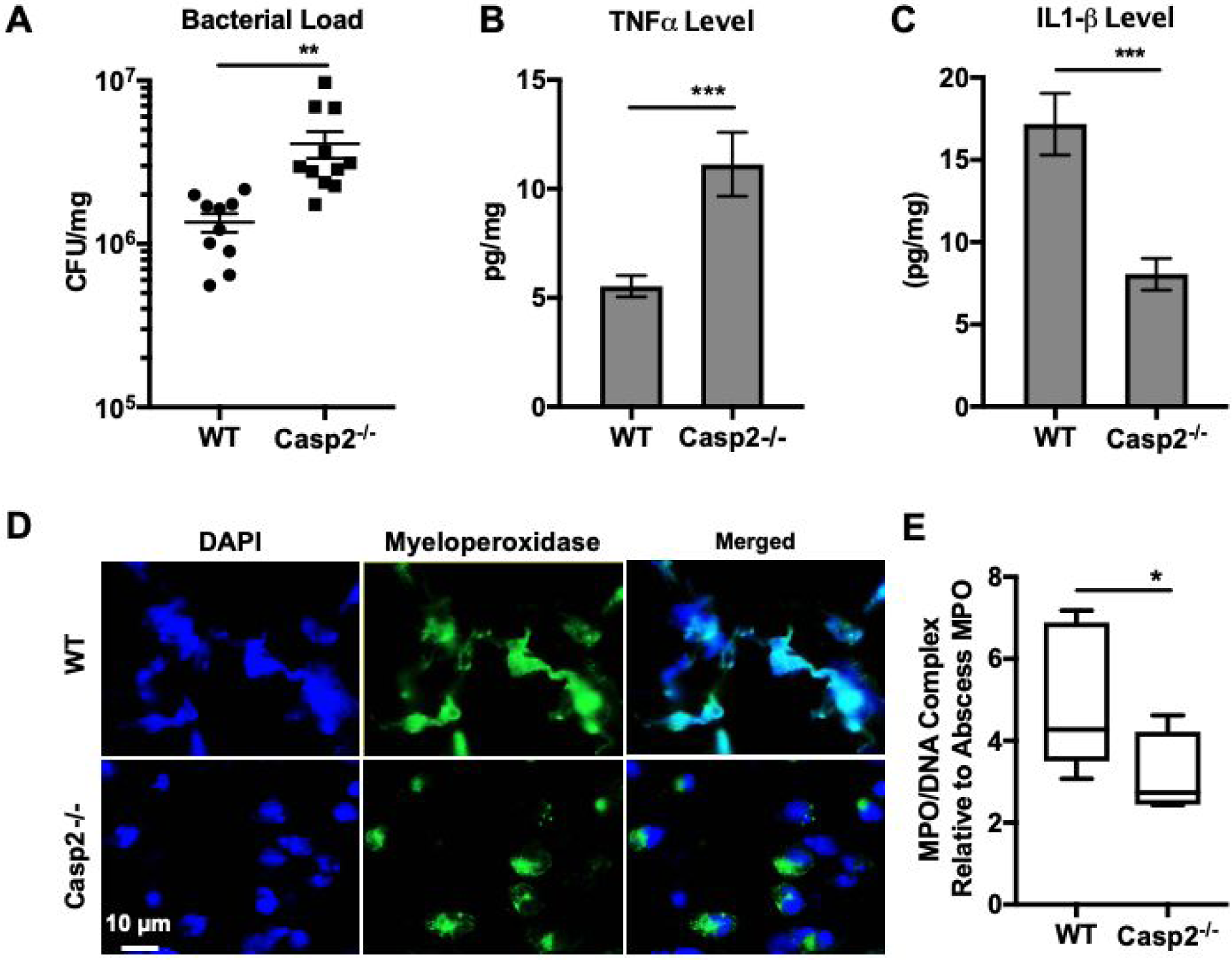
Caspase-2 is required for host defense against MRSA infection and mediates NETosis in the subcutaneous abscess. **(A)** Bacterial burden in abscesses excised from male or female wild-type (WT) and *Casp2^-/-^* C57BL/6 mice infected subcutaneously with 10^7^ CFU MRSA for 3 days. Data are pooled from 2 independent experiments. **(B-C)** TNFα and IL-1β cytokine levels in subcutaneous abscess homogenates of WT and *Casp2^-/-^* mice. Cytokine levels were measured by ELISA. Data are presented as the mean of n=10 WT and n=11 *Casp2^-/-^* mice pooled from 2 independent experiments. **(D)** Representative confocal microscopy images of 0.5 µm histology sections from excised abscess tissue from WT and *Casp2^-/-^* mice infected with MRSA for 2 days. Sections were stained with anti-myeloperoxidase antibody (green) and DAPI (blue) to label DNA. Images were acquired using a Nikon A1 confocal scanning microscope. **(E)** ELISA assay of MPO/DNA complex levels in subcutaneous abscess homogenate from WT and *Casp2^-/-^* mice infected with MRSA for 2 days. Data are shown as box plots with median values of WT (n=4) and *Casp2^-/-^* (n=5) mice. *P* value was calculated using the Mann-Whitney test. *P* value: *< 0.05, **< 0.01 and ***< 0.001.

During *S. aureus* skin infection, NETs are rapidly released by neutrophils to prevent the systemic spread of bacteria (*8*). We next investigated whether NETs could be observed in abscesses in WT and *Casp2^-/-^* mice during MRSA infection, using immunofluorescence microscopy. NET formation was assessed by staining abscess histological sections with DAPI to visualize DNA and for the neutrophil marker myeloperoxidase (MPO) (Fig. 5D). In WT mice, we observed extracellular diffuse DNA associated with MPO, consistent with our observation of NET release in WT neutrophils *ex vivo*. In contrast, extracellular DNA from Casp2^-/-^ mice appeared to be less diffuse and had a multi-lobed appearance. MPO was also localized predominantly within the cells, suggesting that Caspase-2 deficient neutrophils released fewer NETs during MRSA subcutaneous infection. To further quantify NET formation *in vivo*, we used an ELISA assay to measure the level of MPO/DNA complexes, indicative of NET formation, in abscess homogenates of infected mice (Fig. 5E) (*52–54*). MPO/DNA levels were significantly higher in abscesses from WT mice compared to Caspase-2-deficient mice. Collectively, these data suggest that Caspase-2 enhances bacterial clearance by controlling neutrophil NET release *in vivo*.

## Discussion

The IRE1α arm of the ER stress response is integral to development or function of many immune cell types, and has been implicated in many diseases that involve inflammation. Notably, neutrophils can promote inflammatory disease progression and persistence (*34, 56, 57*), but the role of IRE1α in neutrophils during inflammatory challenge had not been well studied. Our previous work identified IRE1α as being required for innate immunity in a subcutaneous model of MRSA infection, where neutrophils predominate (*35, 36*). Our findings here clearly demonstrate that IRE1α is a positive regulator of neutrophil effector function during infection. Activation of IRE1α in neutrophils enhanced production of IL-1β and was required for generation of NETs. We found that inhibition of IRE1α impaired mROS generation, which was needed for CASPASE-2 activation to drive NET formation and IL-1β production *ex-vivo* and in the subcutaneous abscess. Notably, mROS generation was required for NET formation during sterile inflammation, suggesting a common step regulating NET production in response to both infectious and non-infectious stimuli. Lastly, IRE1α mediated MRSA-induced calcium flux independently of mROS production to activate histone citrullination, a key step in the chromatin decondensation that precedes NET formation (*13*). Our work reveals a central role for the ER stress sensor IRE1α in coordinating neutrophil effector functions.

The ER governs many fundamental processes such as protein synthesis, calcium homeostasis and lipid metabolism that occur in all cells. Our data highlight IRE1α as a key regulator of cell type-specific processes as well. For example, IRE1α enhances the ability of dendritic cells (DCs) to process and present antigen to stimulate T cells (*58*). In the liver, IRE1α signaling enhances hepatic lipogenesis, the process by which dietary carbohydrates are converted into fatty acid-forming triglycerides, independently of its role in the ER stress responses (*59*). In macrophages, IRE1α becomes activated via TLR stimulation augmenting macrophage antimicrobial capacity and production of proinflammatory cytokines (*31, 35*). Here, we establish a new role for IRE1α in innate immunity in controlling NET formation, a process that occurs uniquely in neutrophils. These studies emphasize that IRE1α function in particular cell types may encompass both its role in ameliorating ER stress and as an amplifier of signaling, which can impact metabolism and inflammation.

Neutrophils possess lower mitochondrial mass than other cells and mainly rely on glycolysis for ATP production (*60*), thus the contribution of mitochondria to neutrophil function was not historically appreciated. More recent work has demonstrated that neutrophils contain a complex network of mitochondria and that the activity of this organelle augments many essential neutrophil functions including chemotaxis, sustained oxidative stress, NF*κ*B signaling, degranulation and apoptosis (*61–64*). Additionally, mitochondrial products such as mROS and NAD+ produced through the activity of Complex I are required for NET formation during sterile inflammation and in response to other immune signals (*17, 65*). Our data show an essential role for mitochondrial function in the response of neutrophils to infectious challenge. Collectively, our work, together with recent studies, establish mitochondria as a central regulator of neutrophil function.

Calcium release from ER stores serves as a potent stimulatory signal driving immune activation. Neutrophil stimulation with a calcium ionophore increases cytosolic calcium concentration to induce PAD4-dependent histone citrullination and NET formation (*14, 20*). The upstream signals that initiate calcium flux and PAD4 activation in neutrophils have not been well defined, however, our findings now point to neutrophil IRE1α as a mechanism to control calcium influx and histone citrullination in response to bacterial infection. Of note, while IRE1α mediated calcium flux specifically in response to MRSA infection, IRE1α activation was not required for ionophore-dependent calcium flux. Calcium can be released from the ER into the cytosol by the activity of the 1,4,5-triphosphate receptor (InsP3R) channels (*66*), and recent work has implicated InsP3R channels as a mechanism by which IRE1α can mediate flux (*67*). In that case, IRE1α acted as a scaffold independently of its enzymatic activity to concentrate InsP3R channels at ER-mitochondria contact sites to facilitate calcium release. However, our data suggest that IRE1α endonuclease activity is required for calcium flux during MRSA infection, since an inhibitor of the endonuclease domain prevented calcium flux and NETosis. Thus, IRE1α may control calcium flux through multiple mechanisms in a context-dependent manner.

Targeting IRE1α activity with small molecule inhibitors represents a potential therapeutic strategy for treating inflammatory diseases. However, our data show that IRE1α augments host defense against bacterial challenge and its inhibition could inadvertently increase susceptibility to microbial infection (*31, 35*). Some evidence in murine models of inflammatory disease, such as rheumatoid arthritis or atherosclerosis, suggests that IRE1α inhibition can counteract the progression of chronic inflammation (*33, 70, 71*). Because IRE1α contributes to a range of cellular outcomes, such as cell survival, cell death and inflammation, inhibition of its activity could impact different diseases through diverse mechanisms. In an experimental arthritis mouse model, IRE1α inhibition with 4*µ*8C attenuated joint inflammation by decreasing production of proinflammatory cytokines such as IL-1β, IL-6, and TNF-α (*71*). Together, these studies support the investigation of IRE1α inhibitors as possible treatments in human disease. However, our work and others have demonstrated that the IRE1α arm of the unfolded protein response contributes to immunity against bacterial infection (*31, 35*). Thus, further exploration of the specific mechanisms by which IRE1α controls immunity and inflammation will provide needed molecular context for effective therapeutic strategies.

## Materials and Methods

### Ethics Statement

All animals were housed and treated in accordance with an IACUC-approved protocol (PRO00008690) in Unit for Laboratory Animal Medicine (ULAM) facilities at the University of Michigan Medical School. Blood samples were obtained from healthy adult donors according to the protocol approved by the University of Michigan Medical School (HUM00044257). Written consent was obtained from all donors.

### Primary Human Polymorphonuclear Neutrophil Isolation

Blood derived from healthy human volunteers was collected into citrate tubes (Becton Dickinson) and layered into Ficoll-Paque Plus (Sigma-Aldrich). Samples were centrifuged at 1440 rpm in a swinging bucket centrifuge for 20 min at 23°C without braking. Pellets containing red blood cells (RBCs) were allowed to sediment in a 3% Dextran/saline solution (40 min, RT). Supernatants were collected and centrifuged (1600 rpm, 10 min, 4°C). The remaining RBCs in pellets were then lysed with 9 ml sterile water for 20 sec and isotonicity restored by adding 1 ml 10X HBSS. Samples were centrifuged (1600 rpm, 10 min, 4°C) and pellets containing neutrophils re-suspended in PBS. Neutrophils were counted using an Invitrogen automated cell counter and the purity assessed by flow cytometry using FITC anti-CD16 and APC anti-CD15 antibodies (Miltenyi Biotec) as markers characteristic of human neutrophils.

### Bacterial infections

USA300 LAC, a community associated methicillin-resistant *Staphylococcus aureus* strain (MRSA) and its isogenic strain harboring p*SarA*-sGFP plasmid (MRSA-GFP) (Boles and Horswill, 2008) were stored at −80°C in LB medium containing 20% glycerol. Bacteria were streaked onto tryptic soy agar (TSA, Becton Dickinson) plates and selected colonies grown overnight at 37°C with shaking (240 rpm) in liquid tryptic soy broth (TSB, Becton Dickinson). Bacteria were pelleted, washed and re-suspended in PBS. The bacterial inoculum was estimated based on OD_600_ and verified by plating serial dilutions on agar plates to determine colony forming units (CFU). Neutrophils were infected with a multiplicity of infection (MOI) of 10 in RPMI culture medium containing 1% BSA for 4h. For cytokine analysis, samples were treated with Lysostaphin (10 U/ml) to kill extracellular bacteria. Supernatants were collected 24h post infection and cytokines quantified by ELISA at the University of Michigan ELISA core.

### *XBP1* Splicing Assay

Neutrophils were collected by centrifugation and re-suspended in Trizol reagent (Qiagen). Total RNA was extracted by Direct-zol RNA kit according to manufacture protocol (Zymo Research) and quantified by NanoDrop. cDNA synthesis was performed using 500 ng of total RNA, murine leukemia virus reverse transcriptase (RT; Invitrogen), and random hexamers (Applied Biosystems). XBP1 transcripts (spliced and unspliced) were amplified by RT-PCR using PCR master mix (Thermo Fisher) and the following primers: forward (5’-AAACAGAGTAGCAGCTCAGACTGC-3’) and reverse (5’-TCCTTCTGGGTAGACCTCTGGGAG-3’). PCR conditions were: Step 1, 95°C 2 min; Step 2, 35 amplification cycles (95°C 30 sec, 63°C 30 sec, 72°C 30 sec) and Step 3, 72°C 10 min. The PCR product was purified and digested with PstI to discriminate between unspliced (290bp and 183bp) and spliced (473bp) *XBP1*. Spliced and unspliced DNA fragments were resolved by electrophoresis on a 2.5% agarose gel. Band intensities were measured using ImageJ software, and the percent spliced was calculated using the following formula: [*XBP1*_s_/(*XBP1*_s_ + *XBP1*_u_)], which represent the band intensity of the *XBP1* spliced relative to total spliced and unspliced band density.

### Neutrophil Bactericidal Activity

Neutrophils were seeded in 24-well plates at 1 × 10^6^ cells/well in 500 μL of media/well and infected with MRSA (MOI 10). At 4 h, 100 µl aliquot of cell culture media was mixed with 900 μl of (H_2_O_2_ + 0.1% NP-40) to lyse neutrophils and release intracellular bacteria. Subsequently, samples were diluted and plated on TSB plates. CFUs were enumerated on the next day and used to calculate neutrophils percent killing relative to CFU obtained from control sample where bacteria were inoculated without neutrophils.

### Immunofluorescence Microscopy

Neutrophils were first adhered onto poly-lysine coated 1.5mm coverslips in 6-well plates, treated with inhibitors (4μ8C; 25 μM, NecroX-5; 10 μM, Z-VDVAD-FMK; 10 μM) or control DMSO and infected with MRSA-GFP (MOI 10). Cells were fixed at 4h pi with 3.7% paraformaldehyde (overnight, 4°C) and permeabilized with PBS+ 0.1% Triton X-100 for 15 min. Neutrophil Elastase (NE) was visualized using rabbit anti-NE (Abcam ab68672) in staining buffer (PBS, 0.1% Triton X-100, 5% BSA, and 10% normal goat serum). Goat anti-rabbit secondary antibody conjugated to Alexa-594 was used according to manufacturer’s procedure (Thermo Fisher). Coverslipswere mounted on microscope slides using Prolong Diamond (Thermo Fisher). Cells were imaged using an Olympus BX60 microscope. For histology sections, immunofluorescence was performed using rabbit anti-MPO (Dako) followed by a secondary goat anti-rabbit antibody conjugated to Alexa 488 (Thermo Fisher). DAPI was used to stain DNA. Images were taken on the Nikon A1 confocal microscope. All fluorescence images were processed and analyzed using ImageJ.

### SYTOX Green Plate Reader Assay

Neutrophils (2 × 10^4^ cells/well) were seeded onto 96-well plates in the presence of cell-impermeable SYTOX Green DNA-binding dye (500 nM) (Thermo Fisher). Cells were left untreated, treated with control solvent, DMSO or inhibitors (25 μM 4μ8C, 10 μM NecroX-5,10 μM Z-VDVAD-FMK and 10 μM KIRA6). Cells were either left uninfected (Mock) or infected with MRSA (MOI 10). Fluorescence intensity was measured by the Biotek microplate reader with excitation/emission (485/520), at 4h pi. NET formation was normalized to SYTOX Green fluorescence intensity obtained from untreated cell control samples.

### Flow Cytometry Analysis

Total ROS and mitochondrial ROS (hydrogen peroxide), calcium mobilization, mitochondria membrane potential, CASPASE-2 activity and CASPASE-3/7 activity were measured using flow cytometry. For ROS measurements, neutrophils were treated with 10 μM CM-H_2_DCFDA (Thermo Fisher) or 10 μM MitoPY1 (TOCRIS) for 1 hour. Neutrophils were washed twice with media and where indicated, treated for 30 min with inhibitors or control solvent prior to infection with MRSA (MOI 10). For calcium mobilization, neutrophils were loaded with 5 μM Fluo-4 AM (Thermo Fisher) and incubated at 37°C for 30 min. Cells were treated with inhibitors or control DMSO and the baseline fluorescence is acquired prior to addition of calcium ionophore (5 μM A23187) or infected with MRSA (MOI 10). Fluorescence intensity was recorded over time for 300s. Data were analyzed by flowJo software for maximum fluorescence intensity relative to baseline. For other measurements, neutrophils were first treated for 30 min with inhibitors or control solvent and then infected with MRSA (MOI 10) for 4h. Neutrophils were stained with 2 μM JC1 dye (Thermo Fisher) for 20 minutes at 37°C, CASPASE-2 FAM-VDVAD-FMK FLICA substrate (ImmunoChemistry Technologies) for 1h at 37°C or 2 μM CellEvant CASPASE-3/7 Green detection reagent (Thermo Fisher) for 30 minutes at 37°C. Cells stained with CASPASE-2 substrate and JC1 dye were washed twice with PBS prior to flow cytometry analysis. Data were further analyzed with FlowJo software and MFI for each condition was determined as the geometric mean. Ratiometric analysis of red fluorescence (FL2) to green fluorescence (FL1) was used to reflect mitochondrial membrane potential. Percent CASPASE-2^+^ or CASPASE-3/7^+^ cells was determined by gating against mock unstained cells.

### Mouse infection

Subcutaneous MRSA infections were performed as previously described (*73*). Male and Female C57BL/6 mice and *Casp2^-/-^* mice were shaved on the right flank. Mice were inoculated with 10^7^ bacteria in 100 µl of PBS subcutaneously in the shaved area of the skin using a 27 gauge needle. Mice were sacrificed on day 3 pi and abscesses were excised, weighed and homogenized in PBS. Total CFU per mouse abscess was enumerated by serial dilution and plating on TSA agar. Total CFU was converted to CFU/mg of tissue weight. Cytokines were quantified by ELISA at the University of Michigan ELISA core and converted to pg/mg of tissue weight. For histology samples, abscesses were excised on day 2 post infection and fixed in neutral formalin solution. Histology samples were further processed to obtain 5 μm paraffin sections by the Research Histology Core at the University of Michigan Medical School.

### Statistical Analysis

Data were statistically analyzed and graphed using Graphpad Prism 7. Differences between the 2 groups were tested using the Mann-Whitney Test. Differences between 3 or more groups were tested using One-way ANOVA and followed up by Tukey’s multiple comparisons test. The mean of at least 3 independent experiments was presented with error bars showing standard deviation (SD) or standard error of the mean (SEM), as indicated in figure legends. *P* values of less than 0.05 were considered significant and designated by: \**P* < 0.05, \*\**P* < 0.01, \*\*\**P* < 0.001 and \*\*\*\**P* < 0.0001. All statistically significant comparisons within experimental groups are marked.

## Acknowledgements

We thank O’Riordan lab members for many helpful discussions. We thank Dr. Karla Passalacqua for helpful comments and reviewing the manuscript. We gratefully acknowledge the Microscopy and Image Analysis Laboratory (MIL), and the Comprehensive Cancer Center Immunology and Research Histology Cores at the University of Michigan Medical School.

## Funding

This work was supported by NIH awards R21 AI135403 (MXO) and R01 HL134846 (JSK).

## Author contributions

BHA, JSK and MXO designed the experiments; BHA and GJS performed the experiments; BHA and MXO wrote the manuscript; TLS and FG assisted in experimental preparation.

## Conflict of interest

The authors have no competing conflict of interest.

## Data and materials availability

Data used in the study are present in the Figures and Supplementary Material. All reagents used in the study are commercially available. Wild-type and Caspase-2 deficient mice strains were acquired from the Jackson Laboratory.

## References

1. N. Borregaard, J. B. Cowland, Granules of the human neutrophilic polymorphonuclear leukocyte. Blood. 89, 3503–3521 (1997).

2. A. W. Segal, How neutrophils kill microbes. Annu. Rev. Immunol. 23, 197–223 (2005).

3. B. Amulic, C. Cazalet, G. L. Hayes, K. D. Metzler, A. Zychlinsky, Neutrophil function: from mechanisms to disease. Annu. Rev. Immunol. 30, 459–489 (2012).

4. C. Tecchio, A. Micheletti, M. A. Cassatella, Neutrophil-derived cytokines: facts beyond expression. Front. Immunol. 5, 508 (2014).

5. V. Brinkmann et al., Neutrophil extracellular traps kill bacteria. Science. 303, 1532–1535 (2004).

6. N. Branzk, V. Papayannopoulos, Molecular mechanisms regulating NETosis in infection and disease. Semin. Immunopathol. 35, 513–530 (2013).

7. V. Brinkmann, A. Zychlinsky, Beneficial suicide: why neutrophils die to make NETs. Nat. Rev. Microbiol. 5, 577–582 (2007).

8. B. G. Yipp et al., Infection-induced NETosis is a dynamic process involving neutrophil multitasking in vivo. Nat. Med. 18, 1386–1393 (2012).

9. C. F. Urban et al., Neutrophil extracellular traps contain calprotectin, a cytosolic protein complex involved in host defense against Candida albicans. PLoS Pathog. 5, e1000639 (2009).

10. M. J. Kaplan, M. Radic, Neutrophil extracellular traps: double-edged swords of innate immunity. J. Immunol. 189, 2689–2695 (2012).

11. V. Brinkmann, Neutrophil Extracellular Traps in the Second Decade. J. Innate Immun. 10, 414–421 (2018).

12. T. A. Fuchs et al., Novel cell death program leads to neutrophil extracellular traps. J. Cell Biol. 176, 231–241 (2007).

13. Y. Wang et al., Histone hypercitrullination mediates chromatin decondensation and neutrophil extracellular trap formation. J. Cell Biol. 184, 205–213 (2009).

14. D. N. Douda, M. A. Khan, H. Grasemann, N. Palaniyar, SK3 channel and mitochondrial ROS mediate NADPH oxidase-independent NETosis induced by calcium influx. Proc. Natl. Acad. Sci. U. S. A. 112, 2817–2822 (2015).

15. O. Tatsiy, P. P. McDonald, Physiological Stimuli Induce PAD4-Dependent, ROS-Independent NETosis, With Early and Late Events Controlled by Discrete Signaling Pathways. Front. Immunol. 9, 2036 (2018).

16. Q. Remijsen et al., Neutrophil extracellular trap cell death requires both autophagy and superoxide generation. Cell Res. 21, 290–304 (2011).

17. C. Lood et al., Neutrophil extracellular traps enriched in oxidized mitochondrial DNA are interferogenic and contribute to lupus-like disease. Nat. Med. 22, 146–153 (2016).

18. V. Papayannopoulos, K. D. Metzler, A. Hakkim, A. Zychlinsky, Neutrophil elastase and myeloperoxidase regulate the formation of neutrophil extracellular traps. J. Cell Biol. 191, 677–691 (2010).

19. H. D. Lewis et al., Inhibition of PAD4 activity is sufficient to disrupt mouse and human NET formation. Nat. Chem. Biol. 11, 189–191 (2015).

20. P. Li et al., PAD4 is essential for antibacterial innate immunity mediated by neutrophil extracellular traps. J. Exp. Med. 207, 1853–1862 (2010).

21. M. Leshner et al., PAD4 mediated histone hypercitrullination induces heterochromatin decondensation and chromatin unfolding to form neutrophil extracellular trap-like structures. Frontiers in Immunology. 3 (2012), doi:10.3389/fimmu.2012.00307.

22. Y. Wang et al., Human PAD4 regulates histone arginine methylation levels via demethylimination. Science. 306, 279–283 (2004).

23. K. Nakashima, T. Hagiwara, M. Yamada, Nuclear localization of peptidylarginine deiminase V and histone deimination in granulocytes. J. Biol. Chem. 277, 49562–49568 (2002).

24. A. K. Gupta, S. Giaglis, P. Hasler, S. Hahn, Efficient neutrophil extracellular trap induction requires mobilization of both intracellular and extracellular calcium pools and is modulated by cyclosporine A. PLoS One. 9, e97088 (2014).

25. H. Wu, B. S. H. Ng, G. Thibault, Endoplasmic reticulum stress response in yeast and humans. Biosci. Rep. 34 (2014), doi:10.1042/BSR20140058.

26. A. V. Korennykh et al., The unfolded protein response signals through high-order assembly of Ire1. Nature. 457, 687–693 (2009).

27. M. Calfon et al., IRE1 couples endoplasmic reticulum load to secretory capacity by processing the XBP-1 mRNA. Nature. 415 (2002), pp. 92–96.

28. K. Lee et al., IRE1-mediated unconventional mRNA splicing and S2P-mediated ATF6 cleavage merge to regulate XBP1 in signaling the unfolded protein response. Genes Dev. 16, 452–466 (2002).

29. D. Acosta-Alvear et al., XBP1 Controls Diverse Cell Type- and Condition-Specific Transcriptional Regulatory Networks. Molecular Cell. 27 (2007), pp. 53–66.

30. A.-H. Lee, N. N. Iwakoshi, L. H. Glimcher, XBP-1 regulates a subset of endoplasmic reticulum resident chaperone genes in the unfolded protein response. Mol. Cell. Biol. 23, 7448–7459 (2003).

31. F. Martinon, X. Chen, A.-H. Lee, L. H. Glimcher, TLR activation of the transcription factor XBP1 regulates innate immune responses in macrophages. Nat. Immunol. 11, 411–418 (2010).

32. D. N. Bronner et al., Endoplasmic Reticulum Stress Activates the Inflammasome via NLRP3- and Caspase-2-Driven Mitochondrial Damage. Immunity. 43, 451–462 (2015).

33. O. Tufanli et al., Targeting IRE1 with small molecules counteracts progression of atherosclerosis. Proc. Natl. Acad. Sci. U. S. A. 114, E1395–E1404 (2017).

34. R. P. Junjappa, P. Patil, K. R. Bhattarai, H.-R. Kim, H.-J. Chae, IRE1α Implications in Endoplasmic Reticulum Stress-Mediated Development and Pathogenesis of Autoimmune Diseases. Front. Immunol. 9, 1289 (2018).

35. B. H. Abuaita, K. M. Burkholder, B. R. Boles, M. X. O’Riordan, The Endoplasmic Reticulum Stress Sensor Inositol-Requiring Enzyme 1alpha Augments Bacterial Killing through Sustained Oxidant Production. MBio. 6, e00705 (2015).

36. J. S. Cho et al., Neutrophil-derived IL-1βeta is sufficient for abscess formation in immunity against Staphylococcus aureus in mice. PLoS Pathog. 8, e1003047 (2012).

37. W. M. Nauseef, Isolation of human neutrophils from venous blood. Methods Mol. Biol. 1124, 13–18 (2014).

38. B. C. S. Cross et al., The molecular basis for selective inhibition of unconventional mRNA splicing by an IRE1-binding small molecule. Proc. Natl. Acad. Sci. U. S. A. 109, E869–78 (2012).

39. R. Ghosh et al., Allosteric inhibition of the IRE1α RNase preserves cell viability and function during endoplasmic reticulum stress. Cell. 158, 534–548 (2014).

40. I. Mitroulis et al., Neutrophil extracellular trap formation is associated with IL-1β and autophagy-related signaling in gout. PLoS One. 6, e29318 (2011).

41. A. K. Meher et al., Novel Role of IL (Interleukin)-1β in Neutrophil Extracellular Trap Formation and Abdominal Aortic Aneurysms. Arterioscler. Thromb. Vasc. Biol. 38, 843–853 (2018).

42. B. H. Abuaita, T. L. Schultz, M. X. O’Riordan, Mitochondria-derived vesicles deliver antimicrobial payload to control phagosomal bacteria. bioRxiv (2018), p. 277004.

43. B. C. Dickinson, C. J. Chang, A targetable fluorescent probe for imaging hydrogen peroxide in the mitochondria of living cells. J. Am. Chem. Soc. 130, 9638–9639 (2008).

44. M. Oparka et al., Quantifying ROS levels using CM-H2DCFDA and HyPer. Methods. 109, 3–11 (2016).

45. S. Grazioli, J. Pugin, Mitochondrial Damage-Associated Molecular Patterns: From Inflammatory Signaling to Human Diseases. Front. Immunol. 9, 832 (2018).

46. J.-P. Upton et al., Caspase-2 cleavage of BID is a critical apoptotic signal downstream of endoplasmic reticulum stress. Mol. Cell. Biol. 28, 3943–3951 (2008).

47. L. Bouchier-Hayes, The role of caspase-2 in stress-induced apoptosis. J. Cell. Mol. Med. 14, 1212–1224 (2010).

48. D. Azzouz, M. A. Khan, N. Sweezey, N. Palaniyar, Two-in-one: UV radiation simultaneously induces apoptosis and NETosis. Cell Death Discov. 4, 51 (2018).

49. A. S. Rohrbach, D. J. Slade, P. R. Thompson, K. A. Mowen, Activation of PAD4 in NET formation. Front. Immunol. 3, 360 (2012).

50. G. Imre et al., Caspase-2 is an initiator caspase responsible for pore-forming toxin-mediated apoptosis. EMBO J. 31, 2615–2628 (2012).

51. V. Jesenberger, K. J. Procyk, J. Yuan, S. Reipert, M. Baccarini, Salmonella-induced caspase-2 activation in macrophages: a novel mechanism in pathogen-mediated apoptosis. J. Exp. Med. 192, 1035–1046 (2000).

52. H. Kano, M. A. Huq, M. Tsuda, H. Noguchi, N. Takeyama, Sandwich ELISA for Circulating Myeloperoxidase- and Neutrophil Elastase-DNA Complexes Released from Neutrophil Extracellular Traps. Advanced Techniques in Biology & Medicine. 05 (2017), doi:10.4172/2379-1764.1000196.

53. E. Lefrançais, B. Mallavia, H. Zhuo, C. S. Calfee, M. R. Looney, Maladaptive role of neutrophil extracellular traps in pathogen-induced lung injury. JCI Insight. 3 (2018), doi:10.1172/jci.insight.98178.

54. K. Kessenbrock et al., Netting neutrophils in autoimmune small-vessel vasculitis. Nat. Med. 15, 623–625 (2009).

55. A. M. Reimold et al., Plasma cell differentiation requires the transcription factor XBP-1. Nature. 412, 300–307 (2001).

56. C. Rosales, Neutrophil: A Cell with Many Roles in Inflammation or Several Cell Types? Front. Physiol. 9, 113 (2018).

57. A. D. Garg et al., ER stress-induced inflammation: does it aid or impede disease progression? Trends Mol. Med. 18, 589–598 (2012).

58. F. Osorio et al., The unfolded-protein-response sensor IRE-1α regulates the function of CD8α+ dendritic cells. Nat. Immunol. 15, 248–257 (2014).

59. A.-H. Lee, E. F. Scapa, D. E. Cohen, L. H. Glimcher, Regulation of hepatic lipogenesis by the transcription factor XBP1. Science. 320, 1492–1496 (2008).

60. N. Borregaard, T. Herlin, Energy metabolism of human neutrophils during phagocytosis. J. Clin. Invest. 70, 550–557 (1982).

61. G. Fossati et al., The mitochondrial network of human neutrophils: role in chemotaxis, phagocytosis, respiratory burst activation, and commitment to apoptosis. J. Immunol. 170, 1964–1972 (2003).

62. N. A. Maianski et al., Functional characterization of mitochondria in neutrophils: a role restricted to apoptosis. Cell Death Differ. 11, 143–153 (2004).

63. Y. Bao et al., Mitochondria regulate neutrophil activation by generating ATP for autocrine purinergic signaling. J. Biol. Chem. 289, 26794–26803 (2014).

64. J. W. Zmijewski et al., Mitochondrial respiratory complex I regulates neutrophil activation and severity of lung injury. Am. J. Respir. Crit. Care Med. 178, 168–179 (2008).

65. P. Amini et al., Neutrophil extracellular trap formation requires OPA1-dependent glycolytic ATP production. Nat. Commun. 9, 2958 (2018).

66. M. J. Berridge, The Inositol Trisphosphate/Calcium Signaling Pathway in Health and Disease. Physiol. Rev. 96, 1261–1296 (2016).

67. A. Carreras-Sureda et al., Non-canonical function of IRE1α determines mitochondria-associated endoplasmic reticulum composition to control calcium transfer and bioenergetics. Nat. Cell Biol. 21, 755–767 (2019).

68. D. Han et al., IRE1alpha kinase activation modes control alternate endoribonuclease outputs to determine divergent cell fates. Cell. 138, 562–575 (2009).

69. J.-P. Upton et al., IRE1α cleaves select microRNAs during ER stress to derepress translation of proapoptotic Caspase-2. Science. 338, 818–822 (2012).

70. C. Hetz, J. M. Axten, J. B. Patterson, Pharmacological targeting of the unfolded protein response for disease intervention. Nat. Chem. Biol. 15, 764–775 (2019).

71. Q. Qiu et al., Toll-like receptor-mediated IRE1α activation as a therapeutic target for inflammatory arthritis. The EMBO Journal. 32 (2013), pp. 2477–2490.

72. S. E. Logue et al., Inhibition of IRE1 RNase activity modulates the tumor cell secretome and enhances response to chemotherapy. Nat. Commun. 9, 3267 (2018).

73. C. W. Tseng, M. Sanchez-Martinez, A. Arruda, G. Y. Liu, Subcutaneous infection of methicillin resistant Staphylococcus aureus (MRSA). J. Vis. Exp. (2011), doi:10.3791/2528.

