## Supplementary Materials for "The IRE1α stress signaling axis is a key regulator of neutrophil antimicrobial effector function"

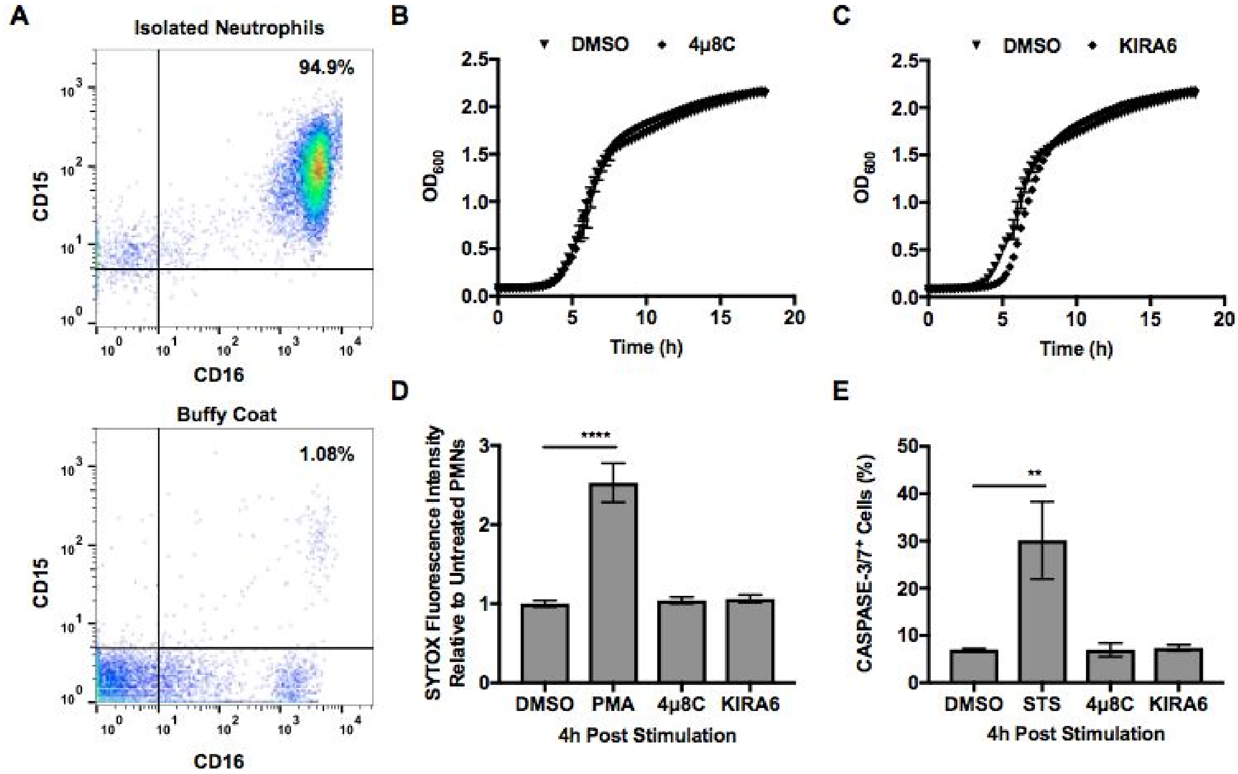


**Fig. S1 IRE1α inhibitors do not impact bacterial and PMN viability.**

**(A)** Isolated human neutrophils and buffy coat were stained with anti-CD15-FITC and anti-CD16-APC and analyzed by flow cytometry. Axenic growth of MRSA in tryptic soy broth in the presence of DMSO or IRE1α inhibitors, 25 μM 4μ8C **(B)** and 10 μM KIRA6 **(C)**. Results represent the average of n≥3 independent experiments done in triplicate. **(D)** SYTOX Green Assay was performed on human neutrophils when left untreated or treated with 100 nM phorbol 12-myristate 13-acetate (PMA), 25 μM 4μ8C and 10 μM KIRA6 and DMSO control for 4h. Graph indicate mean +/- SEM of n≥3 independent experiments. **(E)** CASPASE-3/7 activity of neutrophils treated with DMSO, 1 μM Staurosporine (STS), 25 μM 4μ8C or 10 μM KIRA6 for 4h. CASPASE-3/7 activity was measured by flow cytometry using the CellEvant CASPASE-3/7 green flow cytometry assay kit (Thermo Fisher). Percent of CASPASE-3/7$^{+}$ cells was determined by gating against unstained cells. Graph shows the mean +/- SD from at least 3 independent experiments. *P* value was calculated using one-way ANOVA with Tukey's post-test for multiple comparisons. *P* value: **< 0.01 and ****<0.0001.
